# Engineered RVG29-Conjugated Chitosan for Enhanced Brain Delivery of Kisspeptin Agonists and Antagonists in Reproductive Health

**DOI:** 10.1101/2025.10.26.684692

**Authors:** A Irfan, P Kiran Kumar, Stuti Bhagat, Reena Yadav, Divya Mehta, Anjali Kumari, Aditya D. Deshpande, Bhuvan Bhaskar Tripathi, Y Silambarasan, Mohd Athar, Ananya Aeri, Arvind Sonewane, Vikash Chandra, Sanjay Singh, H.B.D. Prasada Rao, G. Taru Sharma

## Abstract

Efficient delivery of intact therapeutic peptides across the blood–brain barrier (BBB) remains a central obstacle in translating neuroendocrine modulators to clinical practice. Kisspeptin receptor (KISS1R) agonists and antagonists precisely regulate gonadotropin-releasing hormone (GnRH) pulsatility, thereby controlling luteinizing hormone (LH) release, steroidogenesis, and fertility. While the KISS1R agonist KP10 accelerates GnRH–LH pulses to promote ovulation, the antagonist KP234 suppresses pulse generation, inducing reversible infertility. However, their poor BBB permeability limits clinical application. Here, we engineered a non-invasive brain delivery platform by covalently coupling the rabies virus glycoprotein–derived peptide RVG29 to oxidized chitosan nanoparticles via Schiff base chemistry. Sodium periodate oxidation introduced aldehyde groups, FT-IR confirmed aldehyde formation, and UV–Vis spectroscopy validated RVG29 conjugation through characteristic absorbance shifts. The resulting RVG29 - chitosan carriers demonstrated biocompatibility, enhanced neuronal uptake ∼2.1-fold in vitro, and achieved ∼2.5-fold higher brain accumulation in vivo compared to unconjugated controls. Systemic delivery of RVG29–chitosan–KP10 restored and amplified LH pulsatility in ovariectomized mice, accelerated estrous cycling, increased preovulatory follicle numbers, and enhanced litter size. In contrast, RVG29–chitosan–KP234 abolished LH pulses, maintained animals in persistent diestrus, depleted corpora lutea, and produced complete yet reversible infertility. Longitudinal dosing demonstrated stable neuroendocrine modulation without systemic toxicity, and ovarian histology confirmed ovulatory arrest in antagonist-treated animals. By enabling programmable, bidirectional control of a central hypothalamic circuit through systemic delivery of unmodified peptides, this platform addresses a long-standing barrier in neuroendocrine therapeutics. Beyond fertility regulation, this approach can be adapted to other hypothalamic-driven processes, including metabolic, stress-axis, and circadian control.

## Introduction

Peptide therapeutics offer unmatched molecular precision for modulating neuroendocrine circuits, but their application to central nervous system (CNS) disorders is severely limited by the blood–brain barrier (BBB)^1–5^. This tightly regulated endothelial interface effectively excludes most peptides from the brain parenchyma, creating a critical translational gap for hypothalamic-targeted interventions in reproductive disorders, metabolic syndromes, stress-axis dysregulation, and neurodegeneration^1,6–8^. ^9–12^. An ideal delivery strategy would enable non-invasive, systemic administration of unmodified peptides, achieve targeted receptor-mediated transcytosis across the BBB, and precisely engage defined hypothalamic networks^13–15^.

We addressed this challenge by engineering a modular nanocarrier that integrates chitosan, a biodegradable polysaccharide with tunable surface chemistry, and RVG29, a rabies virus glycoprotein-derived peptide that binds nicotinic acetylcholine receptors on BBB endothelial cells to trigger receptor-mediated transcytosis^16–20^. Through covalent conjugation of RVG29 to oxidized chitosan nanoparticles, we generated a BBB-permeable, brain-targeted carrier capable of encapsulating diverse peptide cargos without chemical modification, thereby preserving full bioactivity. This design enables cell-targeted delivery of otherwise BBB-impermeable neuropeptides following systemic injection. As a mechanistic test case, we targeted the kisspeptin–GnRH–LH axis, a fundamental hypothalamic–pituitary–gonadal (HPG) circuit that determines reproductive capacity^21–24^. Kisspeptin, encoded by the KISS1 gene, is produced primarily by KNDy neurons in the arcuate nucleus, which co-express neurokinin B (NKB) and dynorphin. These neurons form a synchronized oscillatory network that generates the pulsatile release of gonadotropin-releasing hormone (GnRH) into the hypophyseal portal system^24–27^. This pulsatility is crucial because rapid GnRH pulses preferentially stimulate luteinizing hormone (LH) secretion, while slower pulses favor follicle-stimulating hormone (FSH) release^28–30^. The frequency and amplitude of these pulses dictate follicular recruitment, ovulation timing, corpus luteum maintenance, and steroid hormone production^29,31,32^. KISS1R agonists such as KP10 increase GnRH pulse frequency and amplitude, leading to elevated LH pulsatility, accelerated follicular development, and enhanced ovulation^33–35^. In contrast, KISS1R antagonists such as KP234 disrupt GnRH pulse generation, suppress LH secretion, and arrest the ovarian cycle in a persistent diestrus-like state, inducing a form of reversible infertility^36–39^. These phenotypes provide highly quantifiable readouts of hypothalamic circuit engagement, making them powerful functional assays for CNS-targeted peptide delivery. In conventional systemic dosing, neither KP10 nor KP234 penetrates the BBB in sufficient quantities to modulate central KNDy neuron activity^40–46^. Effects observed from peripheral administration are largely mediated via pituitary or gonadal pathways, lacking the precision of direct hypothalamic action^42,43,45^. By contrast, delivery via RVG29–chitosan nanocarriers enables intact peptides to cross the BBB, accumulate within the hypothalamus, and engage KNDy neurons directly. This permits true central modulation of reproductive circuitry, driving either sustained enhancement or suppression of LH pulsatility, with downstream effects on estrous cyclicity, follicular dynamics, and fertility outcomes.

Our findings establish that a biomaterial–ligand hybrid nanocarrier can be designed for programmable, bidirectional regulation of a defined hypothalamic circuit after peripheral administration. Beyond reproductive control, this approach provides a broadly applicable framework for delivering peptide therapeutics to hypothalamic networks controlling energy homeostasis, circadian rhythms, stress-axis regulation, and neurodegenerative processes. By combining biomaterials engineering with cell-specific receptor-mediated BBB transport, we address a major translational bottleneck for CNS peptide therapeutics, opening new possibilities for precision neuroendocrine modulation in health and disease.

## Results

### Bioengineering and characterization of RVG29-Conjugated Chitosan for Targeted Peptides Delivery

Chitosan was selected as the delivery scaffold owing to its biocompatibility, biodegradability, and cationic nature, which facilitates electrostatic interaction with negatively charged membranes and enhances cellular uptake^16,47–49^. To enable peptide conjugation, native chitosan was oxidized using sodium periodate (NaIO₄), generating dialdehyde chitosan (DAC) by cleaving vicinal diols at the C2 and C3 positions of glucosamine residues (Fig. 1A, step i). FTIR spectra confirmed aldehyde formation, showing a strong signal at 1770 cm⁻¹ in DAC compared to native chitosan (Fig. 1B). The presence of N–H stretching bands indicated that free amine groups remained available for further functionalization, while consistent CH–OH and CH₂OH signals demonstrated retention of chitosan’s structural integrity. The chemical modification was further verified by Tollens’ test, which produced a silver mirror only in DAC samples, confirming aldehyde group formation (Fig. S1). Next, DAC was conjugated with the brain-targeting peptide RVG29. Using EDC–NHS chemistry, the –COOH group of RVG29 was activated to form a transient O-acylisourea ester, which subsequently reacted with free –NH₂ groups in DAC to yield stable amide linkages (Fig. 1A, steps ii–iii). UV–Vis spectroscopy of purified DAC–RVG29 conjugates displayed a broad absorbance peak between 250–290 nm, distinct from DAC alone and consistent with peptide incorporation (Fig.1C). FTIR spectra further revealed characteristic amide I (1655 cm⁻¹), amide II (1547 cm⁻¹), and amide III (1319 cm⁻¹) bands, together with enhanced C–N stretching at 1050 cm⁻¹, confirming successful peptide conjugation (Fig. 1D). Importantly, HPLC analysis verified the efficiency of conjugation, yielding 85.3 ± 5% for DAC–RVG29 (Fig. S1D–H). The remaining aldehyde groups of DAC–RVG29 were then used to conjugate either kisspeptin-10 (KP10) or kisspeptin-234 (KP234) via Schiff base chemistry (Fig. 1A, step iv). For KP10, UV–Vis spectra showed a red shift in absorbance maxima from 262 nm to 278 nm in purified DAC–RVG29–KP10, consistent with peptide binding (Fig. 1E). No residual KP10 signal was detected in the unconjugated fraction, suggesting complete conjugation. FTIR analysis of DAC–RVG29–KP10 exhibited strong amide I (1658 cm⁻¹) and amide II (1552 cm⁻¹) bands, further supporting successful attachment (Fig. 1F). HPLC confirmed a conjugation efficiency of 82.5 ± 6% for DAC–RVG29–KP10 (Fig. S1D–H). A similar pattern was observed for DAC–RVG29– KP234, with a red shift from 262 nm to 278 nm in UV–Vis spectra (Fig. 1G) and distinct amide I (1663 cm⁻¹), amide II (1544 cm⁻¹), and amide III (1313 cm⁻¹) bands confirming conjugation (Fig. 1H). HPLC analysis verified a conjugation efficiency of 85.5 ± 6% for DAC–RVG29–KP234 (Fig. S1D–H). Together, these results, corroborated by FTIR, UV–Vis spectroscopy, and HPLC, demonstrate the stepwise chemical engineering of chitosan into a multifunctional nanocarrier capable of sequentially conjugating RVG29 for brain targeting and kisspeptin peptides for neuroendocrine modulation.

**Figure 1.**
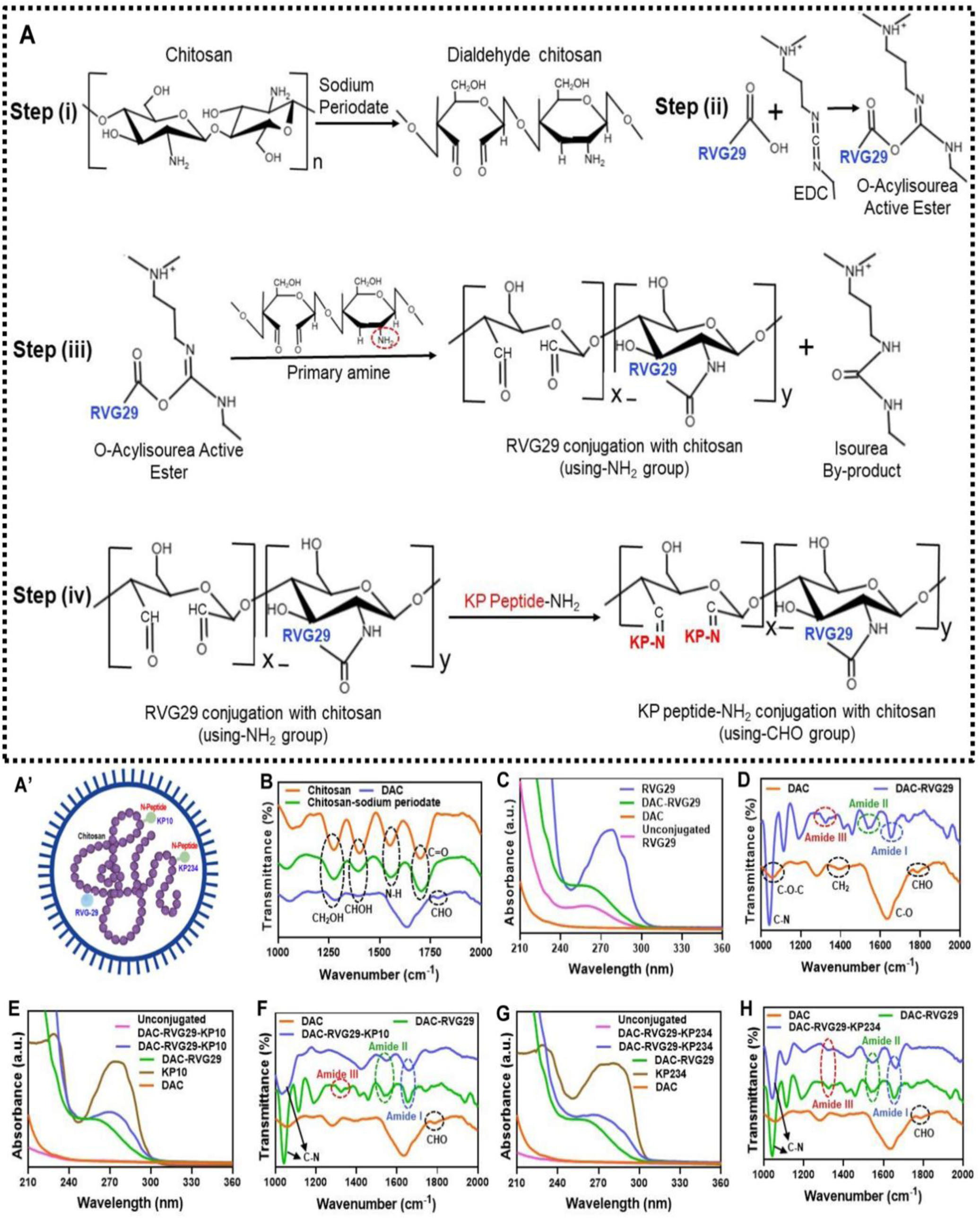
Synthesis and Characterization of Dialdehyde Chitosan Conjugated with Dual Peptides (RVG-KP10/KP234). (A). Schematic representation of the preparation of dialdehyde chitosan (DAC) using sodium periodate followed by conjugation of dual peptide with chitosan. The representative image of DAC-dual peptide formulation is provided in the inset (step-iv). (A’). Schematic representation of Conjugated DAC-RVG-KP10/KP234. (B) FT-IR spectra of chitosan, oxidizing chitosan by sodium periodate and DAC. (C) UV-Vis spectra of free RVG29, DAC-RVG29, DAC and unconjugated RVG29. (D) FT-IR spectra of DAC and DAC-RVG29. (E) UV-Vis spectra of DAC-RVG29, DAC-RVG29-KP10, unconjugated DAC-RVG29-KP10, free KP10 and DAC. (F) FT-IR spectra of DAC, DAC-RVG29 and DAC-RVG29-KP10. (G) UV-Vis spectra of DAC-RVG29, DAC-RVG29-KP234, unconjugated DAC-RVG29-KP234, free KP234 and DAC. (H) FT-IR spectra of DAC, DAC-RVG29 and DAC-RVG29-KP234.

### RVG29 Functionalization Promotes Neuronal Uptake and KISS1R Modulation

To determine whether RVG29 conjugation improved neuronal delivery efficiency, we compared quantum dot (QD)–labelled kisspeptin peptides encapsulated in RVG29–chitosan nanoparticles (RVG29-CS) with QD alone in GT1-7 hypothalamic neurons. High-content imaging revealed intense, punctate intracellular fluorescence in RVG29-CS–treated cells, extending from the cytoplasm into perinuclear regions, consistent with active endosomal trafficking (Fig. 2a,b). Quantitative image analysis confirmed a significant increase in cellular uptake compared with QD controls (p < 0.001), indicating that RVG29 functionalization substantially enhances neuronal entry (Fig. 2c). Cell viability remained above 75% at concentrations up to 50 μg/mL, supporting the biocompatibility of the nanocarrier (Fig. 2d). Functionally, western blot analysis demonstrated that RVG29-DAC–KP234 reduced KISS1R protein abundance significantly, suggesting that targeted antagonist delivery downregulates receptor availability (Fig. 2e).

**Figure 2.**
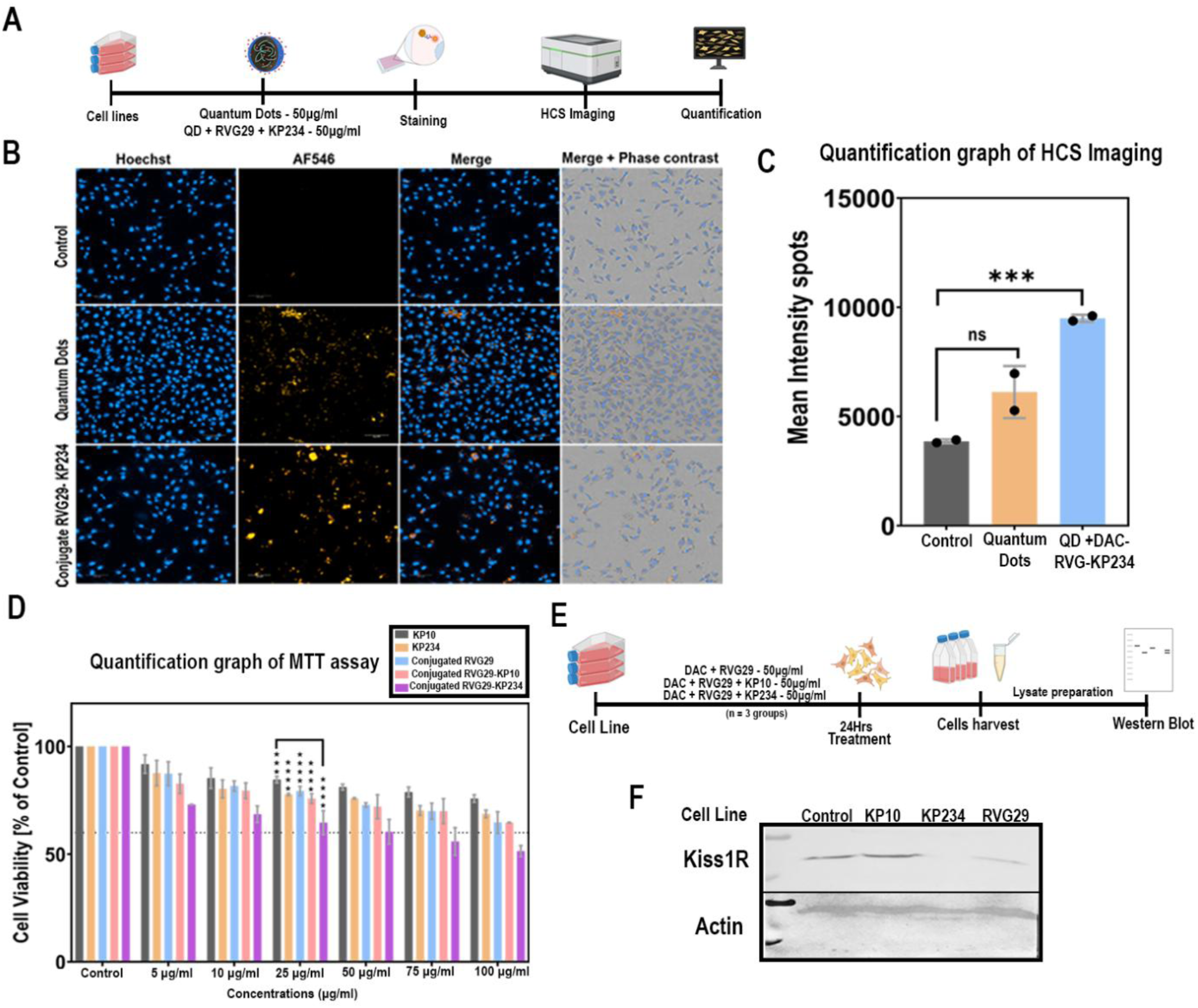
Internalization of Quantum Dots and Peptide-Mediated Modulation of Kiss1r Expression in HPG Cells. (A). Schematic regime using Hpg cell line to check internalization of quantum dots with high content screening image analysis. (B). Hpg cells were stained with nuclei stain (Hoechst 33456), quantum dots at achieving internalization at above 570nm(AF546). (C). Quantification of Hpg cell line internalization graph. Statistical significance at Mean ± SD: using Unpaired t test (**\*\*\***) p=0.003. (D). Quantification of Cell Viability by MTT assay. Statistical significance at Mean ± SD: using Unpaired t test (**\*\*\***), (**\*\*\*\***) p=0.0039, 0.0001. (E). Schematic representation of the experimental workflow in HPG cells treated with conjugated peptides, followed by analysis of protein expression levels using western blotting. (F). Representative western blot image showing Kiss1r protein expression in HPG cells, detected using rabbit anti-Kiss1r antibody. Scale bar = 100um.

### Targeted In Vivo Brain Delivery and Hypothalamic Kisspeptin Signaling Modulation

We next evaluated whether RVG29 functionalization enabled efficient peptide transport across the blood–brain barrier (BBB) and selective engagement of hypothalamic kisspeptin networks in vivo. To establish a traceable system, we used quantum dots (QDs) as model carriers because of their strongfluorescence and stability. Amine-functionalized QDs were conjugated with RVG29 through carbodiimide (EDC–NHS) chemistry, generating stable QD–RVG29 conjugates (Fig.S2A). Subsequent coupling of kisspeptin peptides (KP10 or KP234) to QD–RVG29 produced dual-labeled conjugates, with UV–Vis spectroscopy confirming characteristic blue-shifts in absorbance peaks and FT–IR analysis revealing the presence of amide I (1647 cm⁻¹) and II (1536 cm⁻¹) bonds, validating peptide conjugation (Fig. S2B–C). Importantly, fluorescence emission spectra remained unaltered, demonstrating that optical properties of QDs were preserved after peptide attachment (Fig. S2D). Transmission electron microscopy showed that pristine QDs were spherical and monodisperse (∼6.9 ± 1.14 nm), whereas peptide-conjugated QDs exhibited a slightly reduced size (∼6.3 ± 1.05 nm) and modest clustering, consistent with peptide-mediated surface interactions (Fig. S2E–F).

QD–labeled kisspeptin peptides were administered intravenously in either RVG29–chitosan nanoparticles (RVG29-DAC) or unconjugated QD formulations. Live animal fluorescence imaging revealed rapid brain accumulation in the RVG29-DAC group within 30 min, with strong signal retention for over 4h, whereas unconjugated QDs produced minimal detectable signal (Fig. 3a,b) (Fig. S4a). Ex vivo quantification confirmed a significant increase in brain-associated fluorescence in RVG29-DAC–treated animals (p < 0.001), with limited off-target distribution (Fig. 3c).

**Figure 3.**
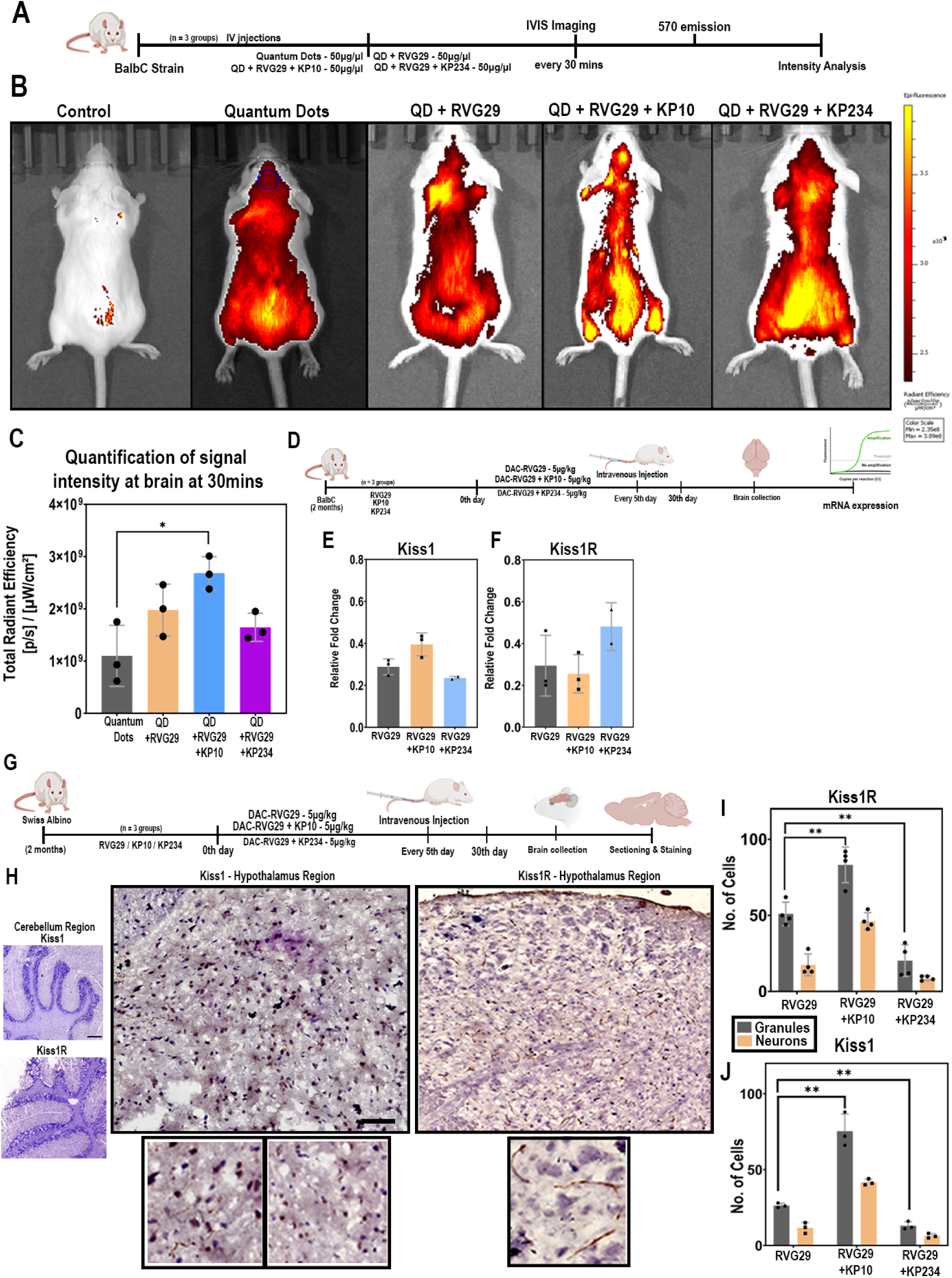
In vivo biodistribution of QD–peptide conjugates, brain gene expression, and protein localization in Balb/cJ mice. (A). Schematic representation of the in vivo imaging system (IVIS) experimental workflow in Balb/cJ mice administered intravenous injections of quantum dots (QDs) conjugated with DAC-conjugated peptides. (B). Representative IVIS images showing comparative distribution of different QD–peptide conjugates in Balb/cJ mice. (C). Quantification of brain signal intensity expressed as total radiance efficiency at 30mins. Data are shown as mean ± SD. Statistical analysis was performed using an unpaired t-test: QD vs. KP10 (p = 0.0051). (D). Schematic representation of the workflow for analyzing mRNA expression signatures in mouse brain tissue using RT-PCR. (E). Quantification of KISS1 gene expression in mouse brain tissue shown as relative fold change. Data are presented as mean ± SD. Statistical analysis was performed using an unpaired t-test: (*) p = 0.0495; (ns) p = 0.1595. (F). Quantification of KISS1R gene expression in mouse brain tissue shown as relative fold change. Data are presented as mean ± SD. Statistical analysis was performed using an unpaired t-test: (ns) p = 0.7153; (ns) p = 0.2279. (G). Schematic representation of the workflow for mouse brain. (H). Representative immunohistochemistry (IHC) images of mouse brain sections showing comparison of the cerebellum and hypothalamic regions. Sections were probed with anti-KISS1 and anti-KISS1R antibodies to visualize protein expression and localization. (I). Quantification of KISS1R expression in cerebellar granular cells and neuronal cell types. Data are presented as mean ± SD. Statistical analysis was performed using an unpaired t-test: () p = 0.003; (*) p = 0.0007; (ns) p = 0.05. (J). Quantification of KISS1 expression in cerebellar granular cells and neuronal cell types. Data are presented as mean ± SD. Statistical analysis was performed using an unpaired t-test: () p = 0.001; (*) p = 0.0002; (ns) p = 0.07.

Molecular profiling of dissected hypothalamic tissue provided evidence of functional engagement: RVG29-DAC–delivered KP10 significantly upregulated Kiss1 mRNA without altering Kiss1r, consistent with agonist-driven transcriptional activation of kisspeptin neurons (Fig. 3d, e). In contrast, RVG29-DAC–KP234 reduced Kiss1 expression while increasing Kiss1r, suggestive of compensatory receptor upregulation under antagonist pressure. Immunohistochemical analysis revealed two distinct kisspeptin and KISS1R staining patterns: dense, granular clusters within the arcuate nucleus and linear neuronal projections extending toward GnRH neuron terminals (Fig. 3g,h). Quantification showed that both patterns were amplified in KP10-treated mice and reduced significantly following KP234treatment (Fig. 3i,j). This bidirectional modulation of discrete hypothalamic kisspeptin microcircuits indicates that RVG29-mediated BBB transport not only achieves central delivery but also allows for targeted rewiring of reproductive neuroendocrine pathways.

### Central Kisspeptin Modulation Bidirectionally Regulated LH Pulsatility and Steroid hormones

Given the central role of kisspeptin in GnRH–LH regulation, we evaluated luteinizing hormone dynamics in ovariectomized females (Fig. 4a). Single-dose KP234 treatment via RVG29-DAC eliminated the subtle, ultradian fluctuations observed in controls (Fig. 4b). High-frequency sampling (every 15 min) demonstrated that KP10 restored ∼30-min LH pulse intervals, whereas KP234 abolished pulsatile release, yielding flat-line profiles (Fig. 4c). In a chronic dosing paradigm (5 μg/kg every 5 days for 30 days), KP234 sustained LH suppression, while KP10 maintained elevated LH pulses relative to RVG29 alone (Fig. 4d,e). Hormone profiling revealed that KP234 dysregulated estradiol, progesterone, and FSH, consistent with a central block in GnRH signaling, whereas KP10 promoted steroid hormone production (Fig. 4f,g,h). These findings confirm that BBB-permeable kisspeptin modulators can precisely and persistently shift neuroendocrine output toward either suppression or stimulation.

**Figure 4.**
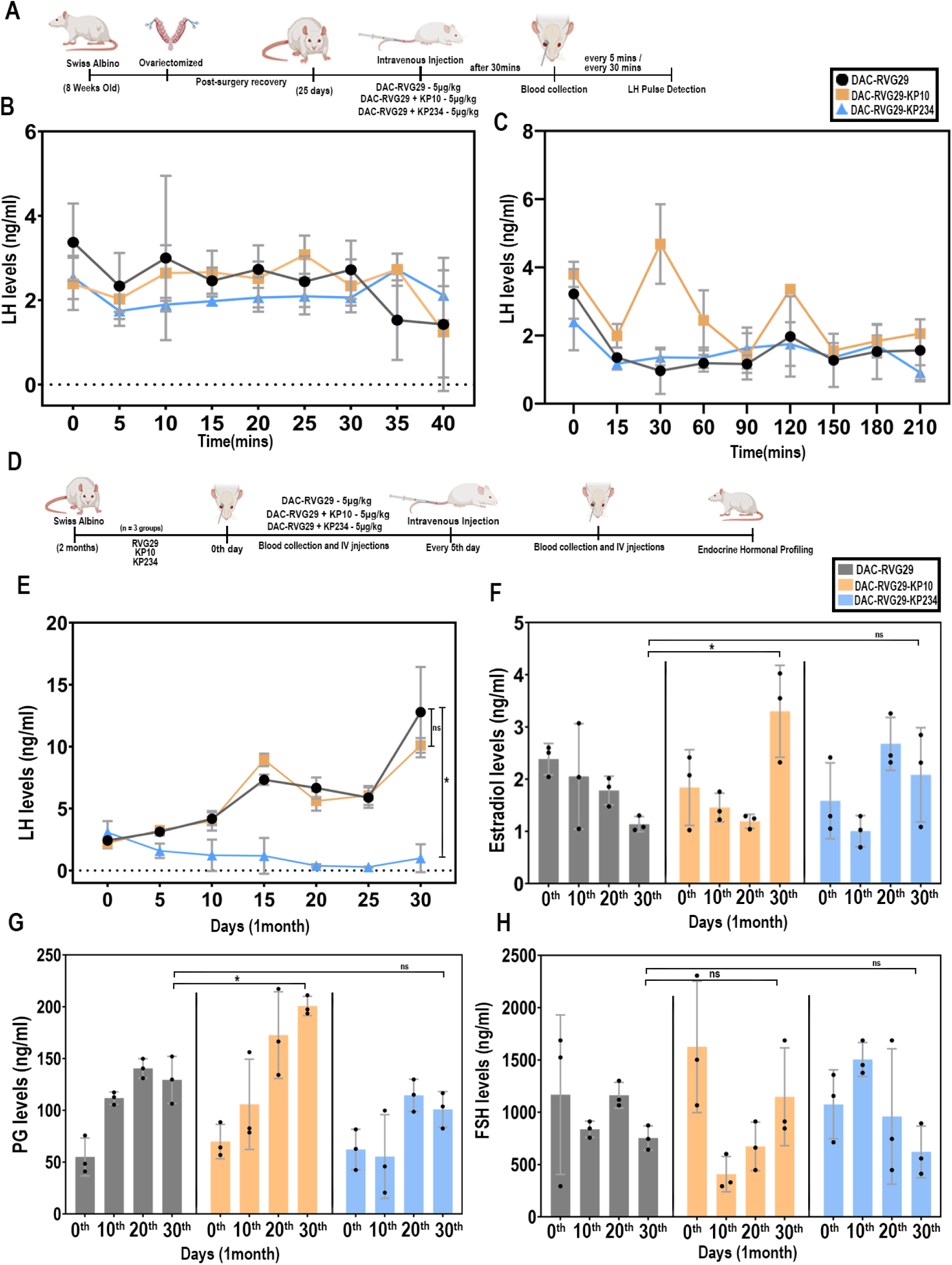
LH pulse and long-term endocrine hormonal profiling with & without ovariectomized mice treated with conjugated peptides. (A). Schematic representation of the experimental workflow in ovariectomized mice for luteinizing hormone (LH) pulse detection. (B). Graph showing luteinizing hormone (LH) levels measured at 5-minute intervals in ovariectomized mice. Data are presented as mean ± SD. Statistical analysis was performed using an unpaired t-test: (ns) p = 0.51; (ns) p = 0.18. (C). Graph showing luteinizing hormone (LH) levels measured at 30-minute intervals in ovariectomized mice. Data are presented as mean ± SD. Statistical analysis was performed using an unpaired t-test: (*) p = 0.01; (ns) p = 0.39. (D). Schematic representation of the experimental workflow in mice treated for 1 month with conjugated peptides, followed by endocrine hormonal profiling. (E). Graph showing luteinizing hormone (LH) levels measured every 5th day over 1 month in mice treated with conjugated peptides. Data are presented as mean ± SD. Statistical analysis was performed using an unpaired t-test: (ns) p = 0.27; (**) p = 0.0059. (F) Graph showing estrogen levels measured every 5th day over 1 month in mice treated with conjugated peptides. Data are presented as mean ± SD. Statistical analysis was performed using two-way ANOVA: (*) p = 0.01; (ns) p = 0.26. (G). Graph showing progesterone (PG) levels measured every 5th day over 1 month in mice treated with conjugated peptides. Data are presented as mean ± SD. Statistical analysis was performed using two-way ANOVA: (*) p = 0.01; (ns) p = 0.39. (H) Graph showing follicle-stimulating hormone (FSH) levels measured every 5th day over 1 month in mice treated with conjugated peptides. Data are presented as mean ± SD. Statistical analysis was performed using two-way ANOVA: (ns) p = 0.68; (ns) p = 0.99.

### Chronic Central Kisspeptin modulation Reshapes Estrous Cyclicity and Fertility Outcomes

To determine whether altered hypothalamic signaling translated into reproductive phenotypes, we monitored estrous cycles during chronic treatment (Fig. 5a). KP234-treated mice remained in a persistent diestrus state, consistent with ovulatory arrest, while RVG29 alone and KP10-treated mice maintained regular cyclicity (Fig. 5a). Fertility trials revealed complete infertility in the KP234 group, whereas KP10 increased litter size relative to controls (Fig. 5b). Ovarian histology linked these outcomes to structural changes: KP234 ovaries showed reduced corpora lutea and accumulated atretic follicles, while KP10-treated ovaries exhibited increased total follicle counts (Fig. 5c,d,e,f). Quantitative analysis confirmed depletion of preovulatory follicles in KP234-treated animals and enhanced early antral recruitment in KP10-treated animals (Fig. 5c,d,e,f). Together, these phenotypes provide a mechanistic bridge from targeted hypothalamic kisspeptin modulation to whole-animal reproductive capacity, demonstrating the platform’s translational potential for controlled fertility regulation.

**Figure 5.**
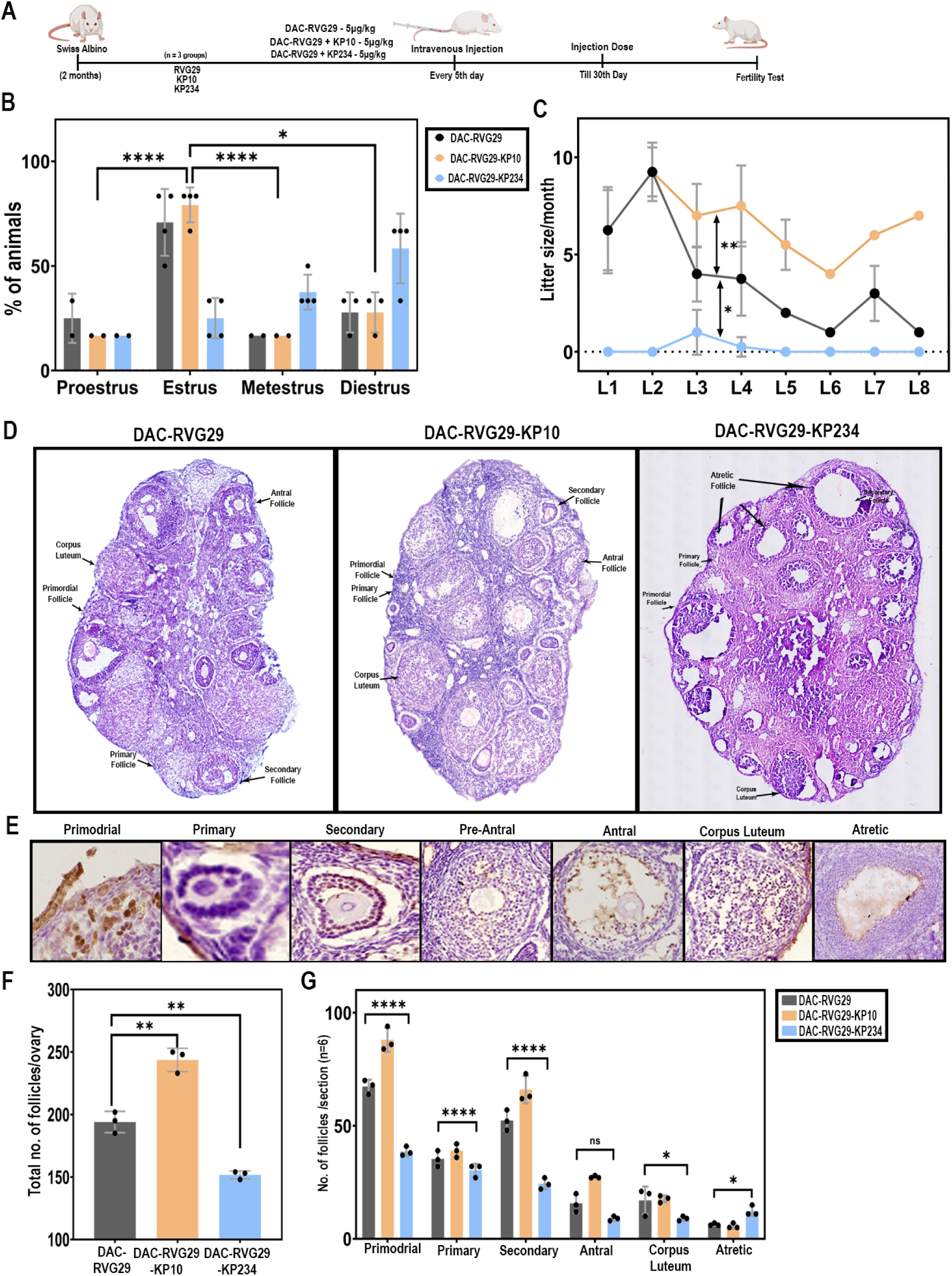
Fertility assessment and ovarian morphology in mice treated with conjugated peptides. (A). Schematic representation of the experimental workflow in mice treated for 1 month with conjugated peptides, followed by fertility testing and estrous cycle monitoring. (B) Graph showing the percentage of animals in different estrous cycle stages averaged over the treatment period. Data are presented as mean ± SD. Statistical analysis was performed using two-way ANOVA: (ns) p = 0.66; (**) p = 0.0021; (****) p = 0.0001. (C) Graph showing litter size per month in mice treated with conjugated peptides. Data are presented as mean ± SD. Statistical analysis was performed using two-way ANOVA: () p = 0.047; () p = 0.0009; (****) p = 0.0001. (D) Representative images of H&E-stained ovaries from mice treated with conjugated peptides. (E) Classification of different ovarian follicle types observed in the study. (F) Quantification of ovarian follicles per ovary. Statistical analysis comparing RVG29 vs. KP10 (p = 0.0024) and RVG29 vs. KP234 (p = 0.0013) using unpaired t-tests. (G) Quantification of follicles per ovarian section. Statistical analysis of atretic follicles comparing RVG29 vs. KP234 using unpaired t-test (p = 0.0120).

## Discussion

### Engineering a BBB Delivery Platform for Peptide Neurotherapeutics

Efficient delivery of bioactive peptides into the central nervous system remains one of the most formidable challenges in translational neuroendocrinology. Systemically administered peptides are rapidly degraded, exhibit poor blood–brain barrier (BBB) penetration, and often require structural modification to achieve therapeutic activity. Such modifications, however, compromise receptor selectivity and complicate clinical translation. Current strategies for bypassing the BBB in peptide therapeutics generally fall into three categories: chemical modification of the peptide cargo to increase lipophilicity or proteolytic stability, nanoparticle encapsulation to improve circulation and uptake, and receptor-mediated transport using Trojan-horse carriers such as transferrin or insulin receptor antibodies. While each has demonstrated proof-of-concept success, their limitations remain significant. Chemical modification alters receptor-binding specificity and often compromises physiological activity, presenting a major translational hurdle in peptide drug development. Encapsulation strategies frequently suffer from poor brain penetration, rapid clearance by the reticuloendothelial system, and uncontrolled release kinetics. Trojan-horse antibody approaches provide higher BBB selectivity but are costly, immunogenic, and difficult to scale.

In contrast, the RVG29-chitosan platform addresses many of these challenges simultaneously. By exploiting RVG29’s innate affinity for neuronal nicotinic acetylcholine receptors, this system enables direct transport across the BBB without requiring chemical modification of the peptide cargo, thus preserving native receptor selectivity and bioactivity. The chitosan backbone offers biocompatibility, biodegradability, and versatile conjugation chemistry, allowing stable yet tunable peptide attachment through Schiff base and amide bonds. Importantly, the platform is inherently modular and scalable: while demonstrated here with kisspeptin agonists (KP10) and antagonists (KP234), the same chemistry could be readily adapted for other neuropeptides of clinical interest, including GnRH analogs for fertility control, oxytocin for social and affective disorders, and corticotropin-releasing hormone (CRH) for stress-related conditions. This scalability, combined with its low-cost synthesis and avoidance of peptide modification, positions the RVG29–chitosan system as a broadly compatible BBB delivery strategy with the potential to expand neuropeptide-based therapeutics well beyond reproductive health.

### Case Study: Kisspeptin Delivery as Proof -of -concept for BBB Targeted Neuropeptides

We selected kisspeptin/KNDy neurons as a test case because hypothalamic kisspeptin neurons, particularly the KNDy subset in the arcuate nucleus, function as the central pacemaker of reproductive physiology by orchestrating gonadotropin-releasing hormone (GnRH) pulsatility. This pulsatile code is not incidental but essential: rapid GnRH pulses preferentially stimulate luteinizing hormone (LH) secretion, whereas slower pulses bias toward follicle-stimulating hormone (FSH). Thus, the tempo of GnRH release dictates whether the reproductive system is primed for follicular recruitment, ovulation, or quiescence. Disruptions in this pulse generator underlie major reproductive pathologies including polycystic ovary syndrome (PCOS), hypothalamic amenorrhea, and premature ovarian insufficiency, underscoring the importance of the KNDy clock as a therapeutic target.

Our RVG29–chitosan delivery system uniquely enables direct hypothalamic engagement by transporting unmodified kisspeptin peptides across the BBB, a feat unattainable with systemic administration alone. Functional outcomes confirm that this strategy provides true central control: delivery of the kisspeptin agonist KP10 enhanced Kiss1 transcription, augmented hypothalamic kisspeptin abundance, and strengthened GnRH neuronal projections, collectively sustaining ovulatory activity and boosting fertility. By contrast, delivery of the kisspeptin antagonist KP234 dampened Kiss1 expression, upregulated Kiss1r as a compensatory response, and suppressed cyclicity, inducing a reversible infertile state. These bidirectional outcomes fertility enhancement with KP10 and contraceptive suppression with KP234 validate the kisspeptin system as a master switch for reproductive control. Importantly, conventional interventions act downstream at the ovary (e.g., gonadotropins, contraceptive steroids) or pituitary, often producing broad systemic effects. In contrast, our approach manipulates the upstream hypothalamic pacemaker itself, offering a level of mechanistic precision that could transform both fertility treatments and non-hormonal contraception. This case study illustrates not only the therapeutic potential of kisspeptin modulation but also the broader utility of the platform. Clinical Translation and Broader Applications.

The successful delivery of kisspeptin peptides across the blood–brain barrier highlights the translational promise of the RVG29–chitosan platform for reproductive medicine. Centrally delivered KP10 could serve as a therapeutic option for patients with hypothalamic hypogonadism, restoring GnRH pulsatility and enabling ovulation in otherwise infertile individuals. It may also find utility in assisted reproductive technologies, where kisspeptin is already being explored as a safer alternative to hCG for triggering ovulation. On the other hand, KP234 delivery represents a novel, centrally acting contraceptive strategy. Unlike conventional hormonal contraceptives that suppress ovarian steroidogenesis and carry systemic side effects, KP234 modulates the hypothalamic pulse generator directly, producing a reversible and non-hormonal state of infertility with potentially fewer off-target consequences. This paradigm shift from peripheral hormone suppression to central pulse regulation could expand contraceptive options while improving tolerability and reversibility.

Beyond human health, the same strategy has broad relevance for veterinary medicine and ecological management. Free-roaming dogs, cats, and wildlife serve as major reservoirs for zoonotic infections such as rabies, toxoplasmosis, and echinococcosis, with unchecked reproduction fueling disease spread. Current methods, such as surgical sterilization, are invasive and difficult to scale, while hormonal contraceptives can be environmentally disruptive. A centrally acting, reversible kisspeptin antagonist delivered via RVG29–chitosan offers a practical, non-surgical solution for population control in companion animals and wildlife. Conversely, KP10 delivery could be harnessed in livestock industries to enhance breeding efficiency, or in conservation settings to improve reproductive outcomes in endangered species. Together, these applications position the RVG29–chitosan kisspeptin delivery system not only as a case study in overcoming the BBB barrier for peptide neurotherapeutics, but also as a versatile tool with bidirectional utility, capable of restoring fertility where it is lost and suppressing fertility where it poses medical, social, or ecological challenges. Finally, the implications of RVG29–chitosan extend beyond fertility regulation. Many neuropeptides and peptide drugs are hampered by poor BBB permeability, limiting their therapeutic scope in neuroendocrine disorders and neurodegenerative diseases. The modularity of this system enabling conjugation of diverse peptides without altering nanoparticle stability or optical properties highlights its potential as a generalizable vehicle for central delivery. By bridgingbiomaterials engineeringwith hypothalamic circuit biology, this work provides a blueprint for precision neuroendocrine therapeutics, with applications spanning reproductive health, veterinary medicine, and beyond.

### Strengths and Limitations

The RVG29–chitosan delivery system offers several distinct advantages for neuroendocrine modulation. By exploiting the nicotinic acetylcholine receptor pathway, RVG29 enables efficient and reproducible transport of kisspeptin peptides across the blood–brain barrier, overcoming a major bottleneck in central peptide therapeutics. Chitosan provides an inherently biocompatible and biodegradable backbone, while its cationic charge enhances cellular uptake, ensuring robust delivery with minimal toxicity. The modular conjugation strategy further permits flexible loading of both agonists (KP10) and antagonists (KP234), thereby enabling either stimulation or suppression of reproductive signalling within the same platform. These features highlight the system’s potential translational breadth, from treating human infertility to managing population control in companion and livestock animals.

However, several limitations remain. Schiff base linkages may be prone to hydrolysis in circulation, potentially leading to premature peptide release before BBB penetration. The possibility of immunogenicity with repeated RVG29 exposure, though low, warrants careful evaluation in chronic dosing regimens. In addition, while conjugation chemistry is precise, scaling the oxidation and peptide-binding steps to GMP-compatible production presents technical challenges. Targeting specificity may also be imperfect, given that RVG29 can engage peripheral cholinergic tissues in addition to neurons. Finally, pharmacokinetics and long-term biodistribution remain incompletely understood, and interspecies variability in BBB transport efficiency could complicate translation from rodent models to larger animals or humans. Taken together, these strengths and limitations underscore both the promise of this approach as a flexible neuroendocrine delivery platform and the critical areas requiring further refinement for clinical or veterinary deployment.

## Materials and Methods

Chitosan (medium molecular weight), EDC (1-ethyl-3-(3-dimethylaminopropyl) carbodiimide), NHS (N-hydroxysuccinimide), MES [2-(N-morpholino)ethanesulfonic acid], sodium periodate, membrane filters (10 kDa), amine-functionalized quantum dots (Em. 580 nm), glucose, and silver nitrate solution were procured from Sigma Aldrich. RVG29, kisspeptin10, and kisspeptin234 were procured from Tocris Bioscience. Sodium chloride was purchased from Himedia. Sodium hydroxide pellets were procured from Puregene. Ammonia solution (30%) was procured from Loba Chemie Pvt. Ltd.

### Preparation of dialdehyde chitosan (DAC)

Chitosan (25 mM) was dissolved in Milli-Q water overnight and subsequently incubated with an equal concentration of sodium periodate solution (NaIO₄, 25 mM) for 3 days until the color turned light orange. After incubation, the solution was dialyzed using a 3.5 kDa dialysis membrane against 0.2 M NaCl (pH 3.0) until the color turned transparent, followed by an additional 3-hour dialysis against deionized water (pH 3.0). The dialdehyde-chitosan dialyzed solution was then filtered through a 0.22μm syringe filter and stored at 4 °C. The presence of aldehyde groups in chitosan was analyzed using attenuated total reflection (ATR) mode in Fourier transform infrared spectroscopy (FT-IR), where 10μL of the sample was loaded on a diamond, followed by monitoring the FT-IR spectra (Nicolet iS50, Thermofisher Scientific).

### Tollens test

0.1 M NaOH was added to 2 mL of 0.1 M AgNO₃ solution to obtain brown precipitate. This precipitate was then dissolved by adding concentrated 30% NH₄OH in a dropwise manner until a clear, transparent solution was obtained. 100 μL of chitosan and DAC swas added to the freshly prepared Tollens’ reagent (2 mL) and the mixture was gently warmed in a water bath at 50-55 °C for 3 hours, followed by monitoring the mirror-like formation in glass test tubes. Glucose was used as a positive control for the Tollens’ test (mirror-like formation).

### Conjugation of RVG29 with DAC

For RVG29 conjugation with DAC, the carboxylic group of RVG29 was activated using EDC and NHS chemistry. In a typical reaction, 500 μg of RVG29 peptide was dissolved in 100 μL of 0.1 M MES buffer (pH: 6.0) followed by addition of 70 μg of EDC and 100 μg of NHS. This suspension was incubated for 30 min at room temperature followed by addition of 500 μL of DAC and further incubated for 1 hour. The DAC-RVG29 formulation was purified using 10 kDa membrane filter and stored at 4 °C until further use. Similarly, carboxyl-activated RVG29 (50 μg) was incubated with - NH₂ functionalized quantum dots (QD, 50 μg) for 3 hours at room temperature to formulate QD-RVG29 and then stored at 4 °C after purification.

### Preparation of DAC-RVG29-Kisspeptin10/Kisspeptin

250 μL of DAC-RVG29 formulation was incubated overnight with 150 μg of kisspeptin10 or kisspeptin234 at 4 °C. The next day, this formulation was purified using 10 kDa membrane filter and stored at 4 °C until further use. Similarly, carboxyl-activated (via EDC-NHS chemistry) kisspeptin10 or kisspeptin234 was incubated with QD-RVG29 complex for 3 hours at room temperature and then stored at 4 °C after350 purification.

### Characterization

The absorbance of the ready formulations: DAC-RVG29, DAC-RVG29 Kisspeptin10/Kisspeptin234, QD-RVG29, QD-RVG29-Kisspeptin10/Kisspeptin234 were recorded using UV-Vis spectra (Take3 plate, Synergy LX, BioTek). The conjugation of ready formulations was confirmed using Fourier Transform Infrared Spectroscopy (FTIR, NICOLET355 iS50 FT-IR, Thermo Scientific) in the range of 525 to 5000 cm⁻¹ equipped with attenuated total reflectance (ATR) mode. Stability of the QD, QD-RVG29, and QD-RVG29-Kisspeptin234 formulation was recorded using multimode plate reader (EnSpire, PerkinElmer) at Ex./Em. 520/580 nm. A total of 100μL volume was utilized to obtain the spectra in dark conditions.

### Animal Experiments

Female BALB/cJ mice (8–12 weeks old; 18–26 g at start) were used. All procedures were performed in accordance with the National Institute of Animal Biotechnology (NIAB) Institutional Animal Ethics Committee (IAEC) guidelines and approved protocols (IAEC permit [IAEC/NIAB/2024/16/HBDPR,2025/19/HBDPR]). Mice were group-housed (2–4/cage) on a 12:12 h light: dark cycle with ad libitum access to food and water. Animals were acclimatized for at least 7 days prior to procedures.

### Ovariectomized mice

Female BALB/cJ mice were used for ovariectomy-based studies investigating luteinizing hormone (LH) regulation. Mice were anesthetized with isoflurane, and bilateral ovariectomy was performed under sterile conditions. Postoperative care included administration of analgesics, routine monitoring to minimize pain and distress, and observation for any surgical complications. Following surgery, mice were housed individually in a temperature-controlled environment with ad libitum access to food and water. Animals were allowed a recovery period of 1 month to ensure complete elimination of endogenous ovarian steroid feedback before initiating LH analysis experiments.

### Peptide conjugate treatment

#### Experimental design

Female BALB/cJ mice were randomly assigned to one of three treatment groups: RVG29 (vehicle control), KP10, or KP234. All peptides were administered intravenously via tail vein at a dose of 5 μg/kg body weight. Prior to experimentation, mice were acclimatized for 10 consecutive days to both handling and the retro-orbital blood collection procedure to minimize stress during sampling. For every protocol described below, a zero-time point pre-bleed was collected before peptide administration.

1. Subtle ultradian LH fluctuations (high-frequency sampling): To investigate acute effects on ultradian LH dynamics, mice received a single IV injection of the assigned treatment. One-hour post-injection, blood samples (∼50μL) were collected from the retro-orbital sinus under brief isoflurane anaesthesia every 5 minutes for 30 minutes. Plasma was separated by centrifugation and stored at – 80 °C until LH quantification.
2. Standard LH pulse profiling: To examine standard LH pulsatility, mice received a single IV injection of the assigned treatment. One-hour post-injection, blood (∼50μL) was collected via retro-orbital sinus under brief isoflurane anaesthesia every 15 minutes for 3 hours. Plasma processing and storage were performed as described above.
3. Sustained modulation of LH pulsatility (chronic dosing): For chronic studies, mice received IV injections of their assigned treatment every 5th day for 30days (days 0, 5, 10, 15, 20, 25, and 30). On each dosing day, a retro-orbital blood sample (∼50μL) was collected prior to injection (pre-bleed) and at specified post-dose time points. Following the final injection on day 30, mice were sacrificed the next day. Ovaries and brain tissues were harvested for downstream analysis. In addition, six mice from the study were retained for fertility assessment via natural mating.

#### Ovary fixation, sectioning and staining

Ovaries were processed accordingly. Freshly collected ovaries were fixed in 10% neutral buffered formalin for 24–48 hours at room temperature, followed by cryoprotection through stepwise immersion in 10%, 20%, and 30% sucrose solutions (w/v in PBS), remaining at each concentration until equilibrated. Tissues were maintained in 30% sucrose for 3 days before sectioning. Cryosections (5μm thickness) were prepared using a Leica CM1950 cryostat and mounted onto positively charged, surface-coated glass slides. For hematoxylin and eosin (H&E) staining, slides were first rinsed in distilled water, immersed in hematoxylin solution for 5 dips, rinsed in tap water, differentiated with one dip in acid alcohol, rinsed again in tap water, and then passed once through 95% ethanol. Slides were then immersed in eosin solution for 1 dip, followed by sequential dehydration in 95% and 100% ethanol for 1 minute each. Clearing was performed in xylene I and II for 15 seconds each, after which slides were dried and mounted with DPX mounting medium. Images were acquired using a Nikon Eclipse Ti2 microscope under brightfield illumination.

#### Brain fixation, sectioning and Immunohistochemistry

Brain tissues were processed accordingly. Freshly collected brains were fixed in 10% neutral buffered formalin for 24–48 hours at room temperature, followed by cryoprotection through stepwise immersion in sucrose gradients of 10%, 20%, and 30% (w/v in PBS), remaining at each concentration until equilibrated. Samples were then maintained in 30% sucrose for 3 days before sectioning. Sagittal cryosections of 20μm thickness were prepared using a Leica CM1950 cryostat and mounted on positively charged, surface-coated glass slides. Immunohistochemistry was performed. Briefly, slides underwent antigen retrieval by incubation at 95 °C for 60 minutes in antigen retrieval buffer (10 mM sodium citrate with 0.05% Tween-20), then cooled to room temperature under running water. Sections were washed twice in distilled water for 5 minutes each, followed by a 1-minute rinse in 1× TBST containing 0.1% Tween-20. Blocking was performed using 10% antibody dilution buffer (ADB) for 15 minutes, repeated twice. Sections were incubated overnight (17 hours) at 4°C in a humid chamber with primary antibodies: Rabbit anti-Kiss1R (PA5-96221, 1:100) and430 Rabbit anti-Kiss1 (PA5-91889, 1:100). After incubation, slides were washed twice in 1× TBST-0.1% Tween-20 for 10 minutes each, blocked again in ADB for 15 minutes (twice), and then incubated with Goat anti-Rabbit Biotin secondary antibody (B-2770, 1:1000) for 1 hour at 37 °C in the dark. Following three washes in 1× TBST-0.1% Tween-20 (10 minutes each), sections were processed using the VECTASTAIN ABC-HRP kit (PK-6100) for 1 hour at 37°C in the dark, rinsed in 1× PBST-0.1% Tween-20 for 5 minutes, and developed with the DAB substrate kit (SK-4100) for 1.5 hours at room temperature in the dark according to the manufacturer’s protocol. Slides were counterstained with hematoxylin, rinsed in distilled water, air-dried, and mounted with DPX mounting medium. Images were acquired using a Nikon Eclipse Ti2 microscope under brightfield illumination.

#### Imaging and Quantification

Immunostained and H&E-stained slides were imaged using a Nikon Eclipse Ti2 microscope equipped with 10×, 20×, and 60× objectives (NA 1.4). Image acquisition was performed under consistent exposure settings, and raw images were processed using NIS-Elements software. For ovarian samples, every fifth section was examined for follicle classification, identifying primordial, primary, secondary, preantral, antral, and atretic follicle. Brain sections were imaged using the same microscopy protocol, focusing specifically on the hypothalamic regions in their entirety. Quantification involved comparing antibody-expressing cells in the hypothalamus to non-expressing regions in the cerebellum for each section. In brain samples, brown DAB staining marked granule and neuronal structures, while hematoxylin counterstaining identified cell nuclei. All quantitative analyses were performed by two independent observers, one of whom was blinded to the treatment groups.

#### Endocrine Hormonal profiling

Blood samples were collected into 1.5 mL centrifuge tubes and allowed to clot at room temperature for 1 hour. Samples were then centrifuged at 6000 rpm for 10 minutes at 4 °C to separate the serum, which was transferred into fresh tubes and stored at −80 °C until further analysis. Serum samples obtained from treatment groups DAC-RVG29, DAC-RVG29 + KP10, and DAC-RVG29 + KP234 were analyzed for hormone profiles, including luteinizing hormone (LH), estrogen, follicle-stimulating hormone (FSH), and progesterone, using commercial ELISA kits: Mouse-LH (KLM0573), Mouse-Estrogen (KLM3348), Mouse-FSH (KLM5595), and Mouse-Progesterone (KLM6101), according to the manufacturers’ instructions.

#### In Vivo Animal Imaging

For in vivo imaging, animals were administered QD or conjugated DAC-peptides via intravenous injection. Imaging was performed every 30 minutes using the IVIS Spectrum system under appropriate anesthesia to minimize movement and physiological fluctuations that could affect image quality. Regions of interest (ROIs) were defined for each image, and quantitative analysis was conducted by measuring the average radiance efficiency to assess the biodistribution and kinetics of the conjugated peptides.

#### Cell Culture

HPG cell lines were utilized throughout the experiment. HPG cells were grown in RPMI-1640 (Gibco-11875085), 10% FBS (Gibco-10270106) and 1× penicillin-streptomycin (Sigma P4333) at 37°C and 5% CO₂. Cells were then treated with conjugate peptides at 50μg/mL and incubated at 37°C and 5% CO₂. The cells were harvested after 24 hours of treatment. Harvested cells were then centrifuged at 15,000 rpm at 4°C for 10 minutes to collect the pellets. The pellets were suspended in lysis buffer (25mM Tris-HCL, pH 8.0; 250 mM NaCl; 1% Triton-X-100; 1% SDS; 2mM MgCl₂; PMSF; Protease inhibitor) followed by sonication and centrifugation at 12,000 rpm, at 4°C for 10 minutes. Supernatants were boiled at 100°C for 5 minutes with 1×Laemmli buffer and loaded in SDS-PAGE to check the protein of interest.

#### Immunocytochemistry

HPG cells were seeded onto 96-well cell culture plates and treated with conjugated peptides with quantum dots (QD) at 50μg/mL, incubated at 37°C and 5% CO₂. After 12 hours, cells were then decanted, washed thrice with 1×PBS and then fixed with 2% Paraformaldehyde (PFA) for 30 minutes at room temperature. Excess PFA was removed using 1× PBS. Hoechst was added for 10 minutes to stain cell nuclei. After 1×PBS wash, these cells were observed in High Content Screening (HCS) to check the internalization and then quantified using cell morphological parameters and intensity calculations with statistical significance performed.

#### MTT assay

Cells were seeded at approximately 10,000 per well in 96-well plates. The following day, cells were treated with KP10, KP234, Conjugated-RVG, Conjugated-KP10, Conjugated-KP234 in respective wells and incubated at 37°C for 24 hours. After incubation, treatments were removed from all wells and 3-(4,5-dimethylthiazol-2-yl)-2,5-diphenyltetrazolium bromide (MTT) solution was added to each well, followed by incubation at 37°C for 3 hours. Then DMSO was added to each well and incubated at 27°C for 15 minutes on a shaker in the dark. After 15-minute incubation, wells were read at absorbance of 570 nm.

#### RNA isolation and mRNA expression

Brain samples collected from 1-month study mice were dissected to sagittal sections; half brains were collected in 1 mL of TRIzol (Cat. No. 9108 Takara) for 250 mg tissue. Samples were then chopped finely, followed by vortexing and centrifugation at 10,000 rpm for 2 minutes at 4°C, then sonicated with pulses of 10 seconds on and 15 seconds off for 6 minutes at 30% amplitude. Samples were then centrifuged at 12,000 rpm for 10 minutes at 4°C. The supernatant was collected and RNA was isolated using the phenol-chloroform method. RNA concentration was measured and cDNA was synthesized according to manufacturer instructions (Cat. No. Takara 6110A). One thousand nanograms of cDNA was used to check the mRNA expression levels of KISS1R, KISS1, PDYN, TAC3, GAPDH in sagittal brain using TB Green SYBR mix (Cat. No. – Takara RR820A) as per manufacturer instructions. The relative expression of the PCR product was quantified using the (−ΔΔCₜ) method following Livak’s approach. Statistical significance was established at p < 0.05.

#### Western Blot

The proteins were separated by SDS-PAGE and then transferred onto a nitrocellulose membrane. The membrane was blocked with 5% skimmed milk for 1 hour at room temperature, followed by overnight incubation at 4°C with primary antibodies. The primary antibodies used were as follows: Rabbit Kiss1R [PA5-96221, 1:100], Rabbit Kiss1 [PA5-91889, 1:100]. After overnight incubation, the blots were washed thrice at 5-minute intervals with 1× TBST containing 0.3% Tween 20 and then incubated with the corresponding secondary antibodies labeled with HRP (Goat anti-Rabbit IgG, 31460) for 2 hours at room temperature. After three rounds of washing with1× TBST containing 0.3% Tween 20 and final wash with 1× TBS, the blots were developed using chemiluminescence. The western blot signals were quantified using ImageJ software.

#### Fertility studies and Vaginal Cytology

Individual adult female mice from treatment groups were housed separately with untreated males to evaluate fertility. If a female failed to conceive within a month, the male partner was replaced. The females underwent plug-checking for a week or until successful mating with a new male, confirmed by the presence of a mating plug. Litter data was generated for an average of 8 litters (L8). The females’ vaginal cytology was performed twice every week for 4 months to evaluate estrous cycle behavior over average months. Using 1×PBS pipetted into the vaginal opening, the precipitate was collected into 1.5 mL tubes. Droplets of duplicates were placed onto positively charged surface-coated slides and air-dried, followed by 0.1% Trypan blue staining for 1 minute and two washes with dH₂O for 30 seconds each. The stages of estrous cycles were then evaluated and statistical significance was established by comparing different estrous cycle phases over average months.

#### Primer sequences

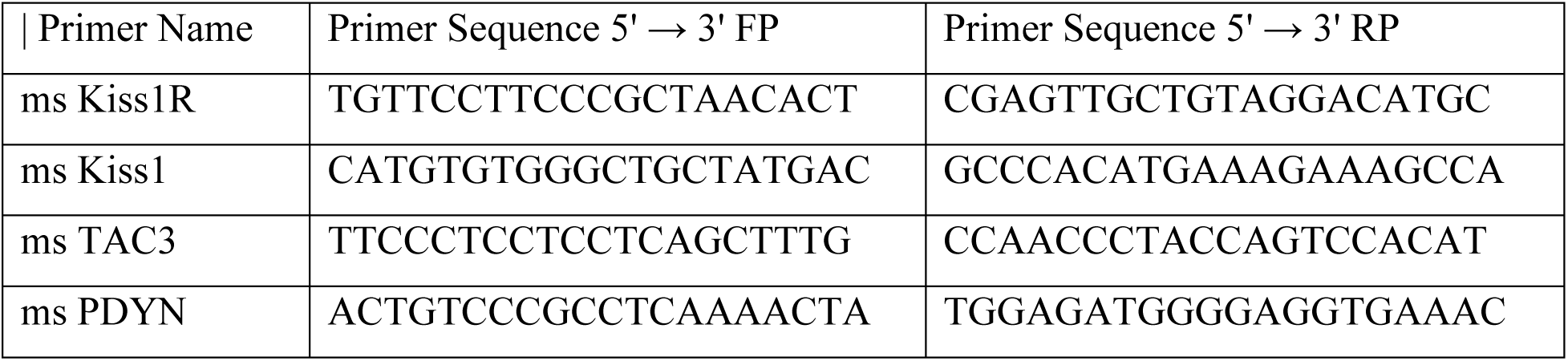

#### Statistical Analysis

Data are presented as mean ± standard error of the mean (SEM). Statistical analyses were performed using GraphPad Prism software. Comparisons between two groups were analyzed using unpaired Student’s t-test, while multiple group comparisons were performed using one-way ANOVA followed by Tukey’s post-hoc test. For time-course experiments, two-way ANOVA with Bonferroni correction was used. Statistical significance was set at p < 0.05 (), p < 0.01 (), and p < 0.001 ().

## Data Availability

The datasets generated and analyzed during the current study are available from the corresponding author upon request. Raw imaging data, hormone profiles, and molecular analysis results supporting the conclusions of this article are included within the manuscript and its supplementary information files.

## Acknowledgments

We thank the National Institute of Animal Biotechnology (NIAB) animal facility staff for excellent animal care and the institutional imaging facility for technical support. We acknowledge the contributions of laboratory members who assisted with tissue processing and data analysis. This work was supported by grants from the NIAB core to H.B.D.P.R., S.S., and G.T.S.

## Competing Interests

The authors declare that they have no competing financial interests or personal relationships that could have appeared to influence the work reported in this paper.

## Supplementary Information

**Figure S1:**
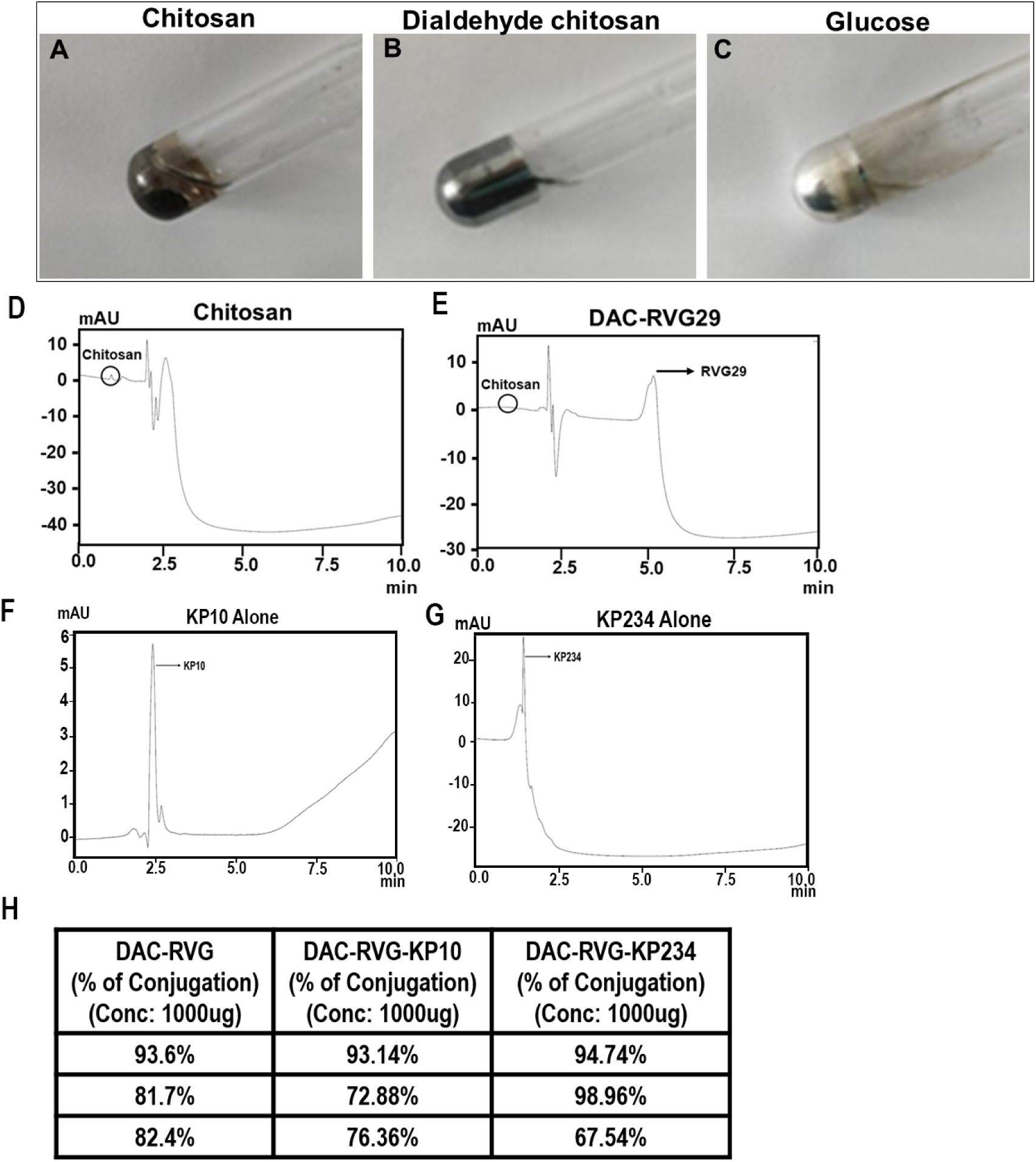
(A-C). Pictures showed Tollens’ test in presence of chitosan (A) and DAC (B) and glucose (C). Tollens’ test validation of aldehyde formation in dialdehyde chitosan (DAC). Silver mirror formation confirms successful oxidation of chitosan to introduce aldehyde groups necessary for peptide conjugation. (D). Spectral image of HPLC showing chitosan peak. (E). Spectral image of HPLC showing DAC-RVG29 peak. (F). Spectral image of HPLC showing KP10-Alone peak. (G). Spectral image of HPLC showing KP234-Alone peak. (H). Table showing concentration of peptide conjugated with RVG29, KP10, KP234 comparing AT 1000ug concentration.

**Figure S2:**
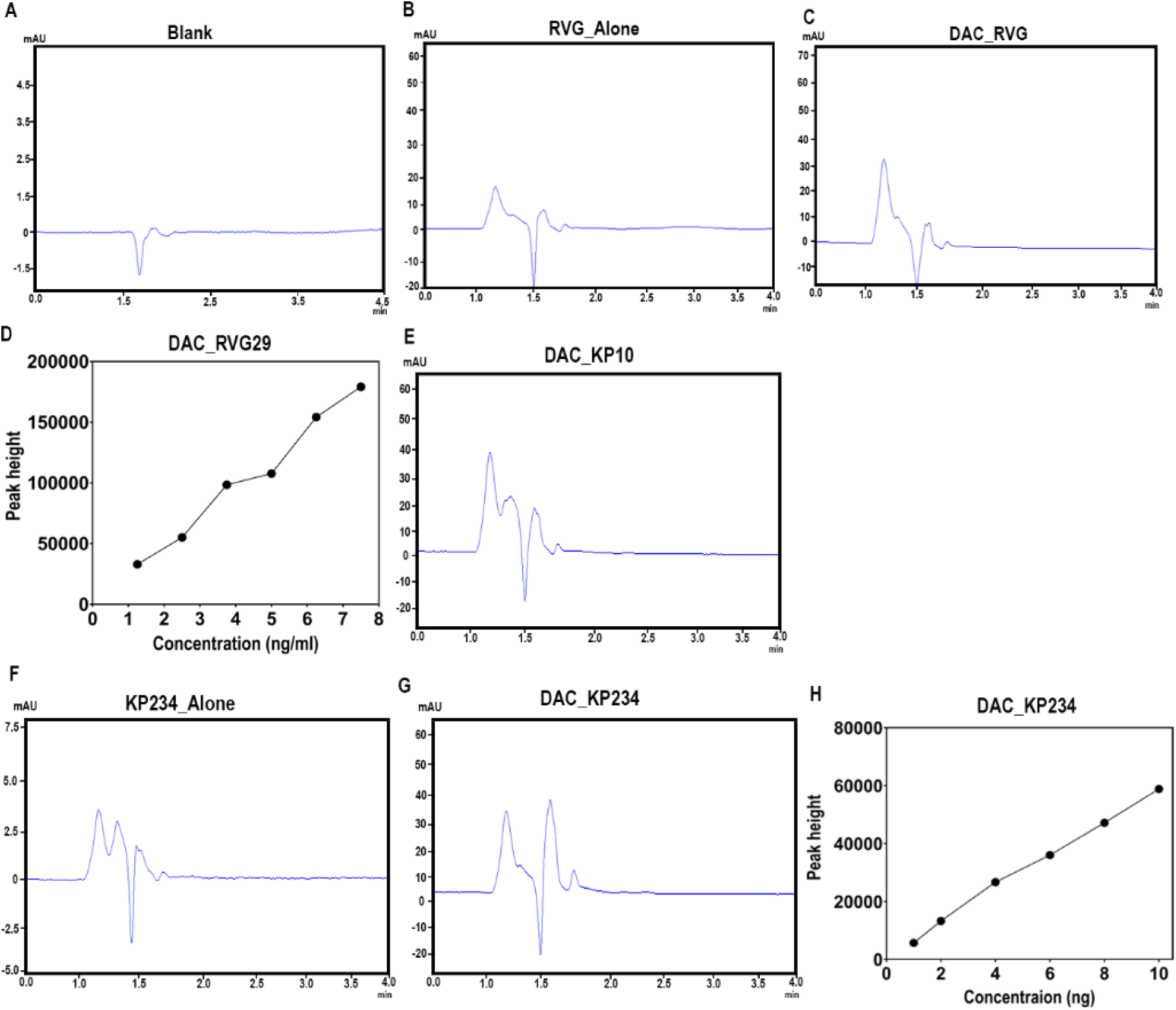
Characterization of peptide conjugates. (A)HPLC spectral image of blank reading at 240nm. (B). HPLC spectral image of RVG-Alone reading at 240nm. (C). HPLC spectral image of DAC-RVG29 reading at 240nm. (D). Standard concentration of RVG-Alone peptide with increasing in concentration. (E). HPLC spectral image of DAC-RVG29-KP10 reading at 240nm. (F). HPLC spectral image of KP234 reading at 240nm. (G). HPLC spectral image of DAC-RVG29-KP234 reading at 240nm. (H). Standard concentration of KP234-Alone peptide with increasing in concentration.

**Figure S3:**
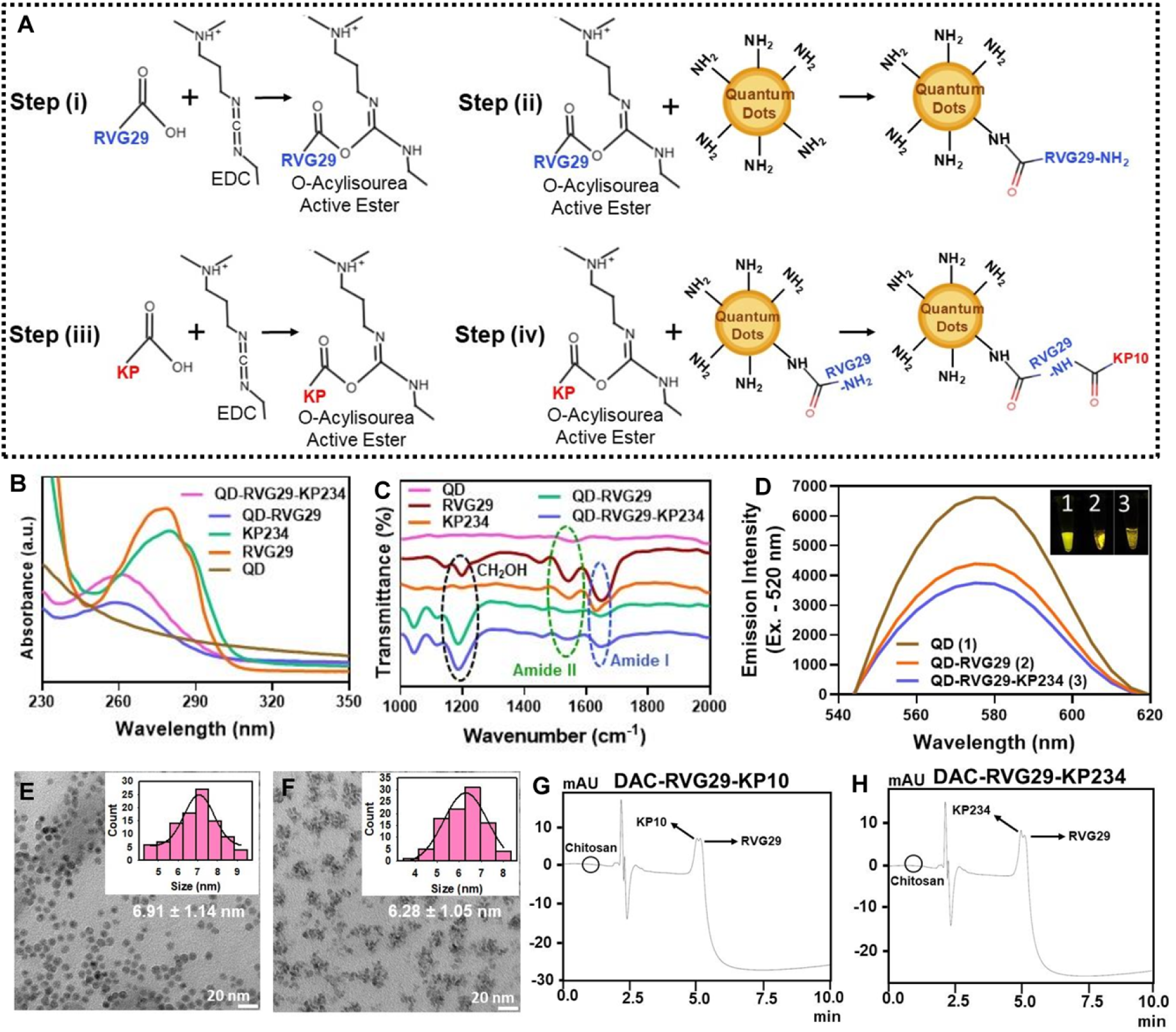
Schematic representation of the activation of RVG29 followed by conjugation of dual peptide with QD (A). FT-IR spectra of QD, free RVG29, free KP234, QD-RVG29 and QD-RVG29-KP234 (B). FT-IR spectra of QD, free RVG29, free KP234, QD-RVG29 and QD-RVG29-KP234 (C). Emission spectra of free QD, QD-RVG29 and QD-RVG29-KP234 with yellow color emission upon ultraviolet light (inset) (D). TEM image of QD (E) and QD-RVG29-KP234 (F). HPLC spectra of DAC-RVG29-KP10 (G) and DAC-RVG29-KP234 formulation (H).

**Figure S4:**
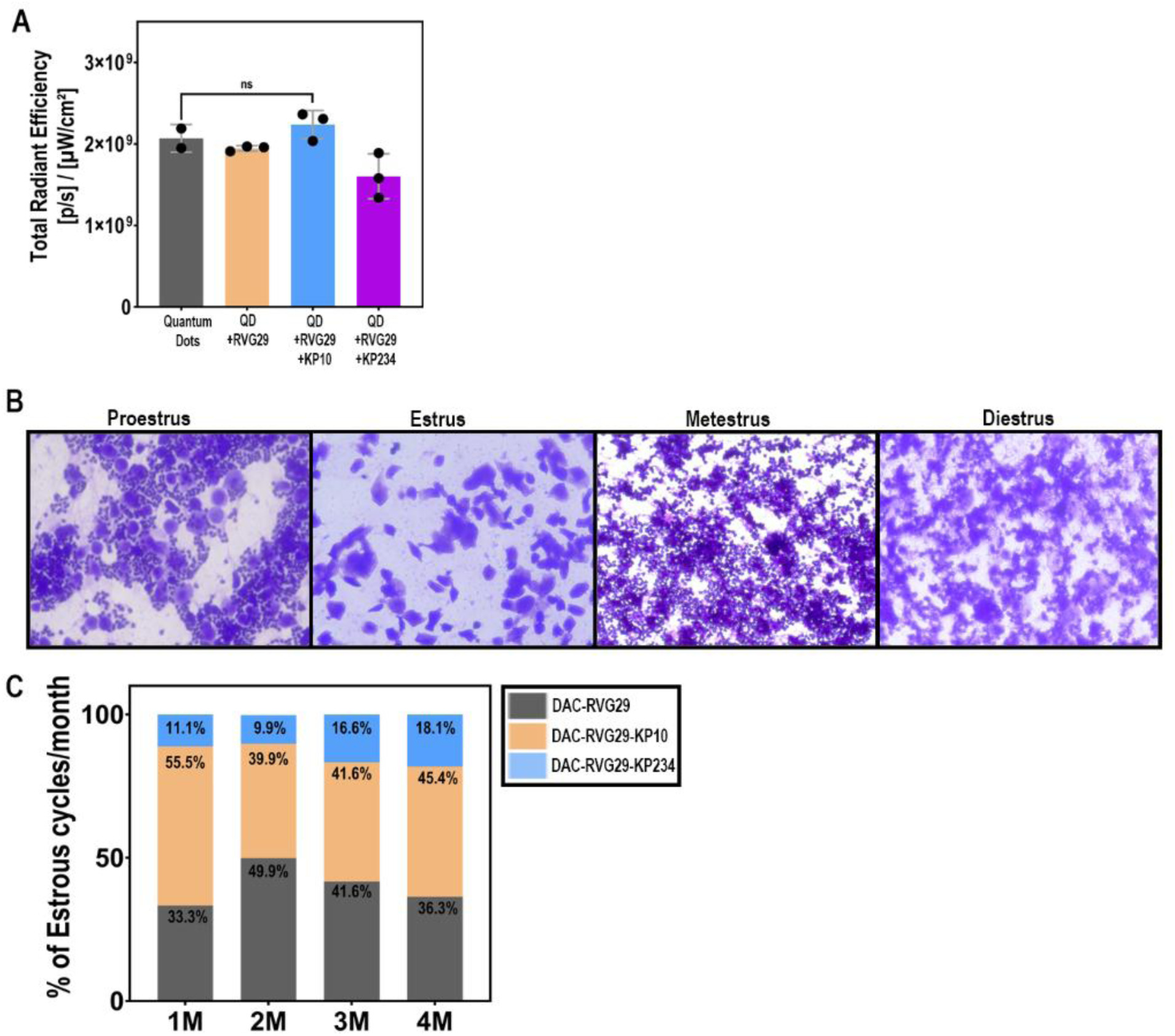
(A). Quantification of brain signal intensity expressed as total radiance efficiency at 4hrs. Data are shown as mean ± SD. Statistical analysis was performed using an unpaired t-test: QD vs. KP10 (p = 0.0051). (B). Representative image of Proestrus, Estrus, Metestrus, Diestrus stained with 0.1% trypan blue. Scale=100um. (C). Percentage distribution of estrus cycles per month among DAC-RVG29, DAC-RVG29-KP10, and DAC-RVG29-KP234 groups.

## Extended Materials and Methods

### Nanoparticle Size and Zeta Potential Analysis

Dynamic light scattering (DLS) measurements were performed using a Malvern Zetasizer Nano ZS to determine the hydrodynamic diameter and polydispersity index (PDI) of chitosan nanoparticles before and after peptide conjugation. Zeta potential was measured in deionized water at pH 7.0 to assess surface charge characteristics that influence cellular uptake and stability.

### Peptide Conjugation Efficiency

The efficiency of peptide conjugation to chitosan carriers was quantified using high-performance liquid chromatography (HPLC). Unconjugated peptides were separated from conjugated formulations using molecular weight cutoff filters, and the concentration of free peptide in the filtrate was measured using UV detection at 280 nm. Conjugation efficiency was calculated as: (Initial peptide concentration - Free peptide concentration) / Initial peptide concentration × 100%.

### Ethical Considerations and Regulatory Compliance

All animal procedures were conducted in accordance with the Guide for the Care and Use of Laboratory Animals and approved by the Institutional Animal Care and Use Committee (IACUC). Animal welfare was monitored continuously, with established endpoints for minimizing pain and distress. Environmental enrichment was provided to enhance animal well-being, and all personnel involved in animal care received appropriate training and certification.

### Quality Control and Reproducibility

Experimental protocols included appropriate controls at each step, including negative controls, vehicle controls, and positive controls where applicable. Critical experiments were repeated by multiple investigators to ensure reproducibility. Standard operating procedures (SOPs) were established for all key methodologies, and regular calibration of analytical instruments was performed to maintain data quality and consistency.

## Notes

### Competing Interest Statement

The authors have declared no competing interest.

